# Patterns of extreme outlier RNA expression in population data reveal sporadic over-activation of genes with co-regulated modules in subsets of individuals

**DOI:** 10.1101/2024.10.04.616600

**Authors:** Chen Xie, Sven Künzel, Wenyu Zhang, Cassandra A. Hathaway, Shelley S. Tworoger, Diethard Tautz

**Affiliations:** Department of Evolutionary Genetics, Max-Planck Institute for Evolutionary Biology, Plön, Germany; Biomedical Pioneering Innovation Center, Peking University, Beijing 100871, China; Shaanxi Key Laboratory of Qinling Ecological Intelligent Monitoring and Protection, School of Ecology and Environment, Northwestern Polytechnical University, Xi’an 710129, China; Research & Development Institute of Northwestern Polytechnical University in Shenzhen, Shenzhen 518063, China; Department of Cancer Epidemiology, Moffitt Cancer Center, Florida, USA; Division of Oncological Science, Knight Cancer Institute, Oregon Health and Science University, USA

**Keywords:** transcriptome analysis, outlier expression, mice, humans, Drosophila, family study

## Abstract

**Background:** Most RNA-Seq datasets harbor genes with extreme expression levels in some samples. Such extreme outliers are usually treated as technical errors and are removed from the data before further statistical analysis. Here we focus on the patterns of such outlier expression.

**Results:** Our study is based on multiple datasets, including outbred and inbred mice, humans from the GTEx dataset, different *Drosophila* species and single-nuclei sequencing experiments from human brain tissues. All show comparable general patterns of outlier expression. Different individuals can harbor very different numbers of outliers, with some individuals showing extreme numbers in only one out of several organs of the respective individual. A three-generation family analysis in mice was generated and analyzed for the inheritance of outlier patterns. We find that most extreme over-expressions are not inherited, i.e., appear to be sporadically generated in individuals. Still, co-regulated outlier expressions are identifiable in various gene groups, and some correspond with known pathways. Among the co-regulated genes with extreme outlier expression are also the hormone genes prolactin and growth hormone, both in mice and humans, for which we include also protein level data from human cohorts.

**Conclusions:** We show that outlier patterns of gene expression are a biological reality occurring universally across tissues and species. Most of the outlier expressions are spontaneous and not inherited. We discuss the possibility that the outlier patterns reflect edge of chaos effects that are expected for systems of non-linear interactions and feedback loops, such as gene regulatory networks.

## Background

Studying gene expression variation through transcriptome analysis has become the standard in unraveling genetic networks, studying cell differentiation and development, exploring patterns of molecular evolution as well as getting insights into genetic diseases. Expression levels are determined through a statistical sampling process of the number of reads obtained in a high-throughput sequencing experiment. Significant differences in expression are estimated based on variance estimates between replicates (1, 2). Variances should ideally follow a particular statistical model, e.g., a Poisson distribution, to properly calculate error probabilities and corrections. However, real data from RNA-Seq experiments usually exhibit overdispersion, which means that the variance of the observed counts is larger than the mean, sometimes due to extreme outlier expression in single or few biological replicates.

Overdispersion is usually ascribed to various factors, including technical noise, biological variability, or measurement error. It is therefore common practice to log-transform the primary data, since this leads to a down-weighting of these factors. Since overdispersion affects also statistical power for differential expression analysis, standard analysis programs, such as DESeq2, edgeR, or limma-voom (1, 3, 4), use a negative binomial model with dispersion estimation and adjust the variance of the counts. For most genes this helps to account for the overdispersion and to improve the accuracy and robustness of the differential expression analysis.

However, it is known that some genes can show extreme expression values in some individuals, which cannot be corrected with overall statistical analyses. Part of the general recommendation for expression data analysis is therefore the removal of such outlier individuals. A standard practice is to identify individuals with extreme expression values, for example in a PCA (5) or a specific denoising pipeline (6, 7). Such samples are then usually discarded without any further analysis.

These procedures are statistically justifiable under the assumption that outlier values are due to technical error, a concern that was particularly relevant when expression levels were measured via microarray technology. However, with the advent of highly standardized sequencing protocols, the probability of technical error became negligible, obviating even the need to generate technical replicates for quality control (8). Hence, it is possible that outlier values may have a biological meaning that is underexplored because of standardized pre-filtering steps.

A first dedicated analysis of rare outlier expression was done in human data (the GTEx resource: www.gtexportal.org). It aimed to identify rare genetic variants with possible major effects in individuals (9). The algorithm to identify these included the condition that the same outlier expression should occur in at least five tissues to ensure that it was actually a genetic polymorphism that had caused them. Many such cases were detected, but many more were again discarded, since they did not conform to the filter criterium.

In the present study, we analyze several large transcriptome datasets from mice, humans and *Drosophila*, focusing on outlier expression values, even if they occur only in single or few individuals and in single tissues. We show for a sub-dataset that the outliers are fully reproducible in independent sequencing experiments, i.e., should not be considered as technical noise. By using a three-generation family analysis in mice, we show that most of the outlier expression effects are not genetically inherited. We conclude that outlier expression is a biological reality that may be linked to chaos effects that cause sporadic over-activation of transcription of different sets of genes in different individuals.

## Results

The study is based on normalized transcript fragment count data (transcripts per million – TPM for mice and humans, counts per million – CPM for *Drosophila*, and normalized counts for snRNA-seq of humans) from population samples. Since we were specifically interested in identifying outlier patterns, we did not log-transform the data and did not exclude any individuals. We used datasets from mice, humans and *Drosophila* (all sample lists in suppl. Table S1).

The mouse dataset includes 48 individuals derived from an outbred stock of *M. m. domesticus* originally collected from France (DOM), as well as between 19 to 20 individuals from outbred stocks of *M. m. musculus* (MUS), *M. spretus* (SPR) and *M. spicilegus* (SPI). Transcriptome data were derived for five organs from each mouse (10). Further we used brain transcriptome data from 24 individuals from the mouse inbred strain C57BL/6 (suppl. Data S1A-S1E).

The human dataset includes 51 individuals for which three organs overlapping with the mouse organs were available in the GTEx dataset (11) plus a subset of 40 of these individuals from which pituitary data were available (suppl. Data S1F).

The *Drosophila* dataset included data from two species. The first is from *D. melanogaster*, comprising 27 individuals, for which transcriptomes were separately obtained for head and trunk (12). The second set included four population samples from *D. simulans*, with 19 to 22 individuals each and transcriptomes obtained from whole flies (13) (suppl. Data S1G).

### Identification of outliers

In RNA-Seq data, the expression variance for many genes conform to a random distribution model. This can be visualized in quantile-quantile (Q-Q) plots that compare the actual distribution of data points with a normal distribution modelled on these datapoints. When the data conform to a normal distribution, one will see a straight line (Figure 1A, left two plots). However, for some genes, there are outlier expression values in some individuals, far away from the line, not compatible with a random distribution (Figure 1A, right two plots).

**Figure 1:**
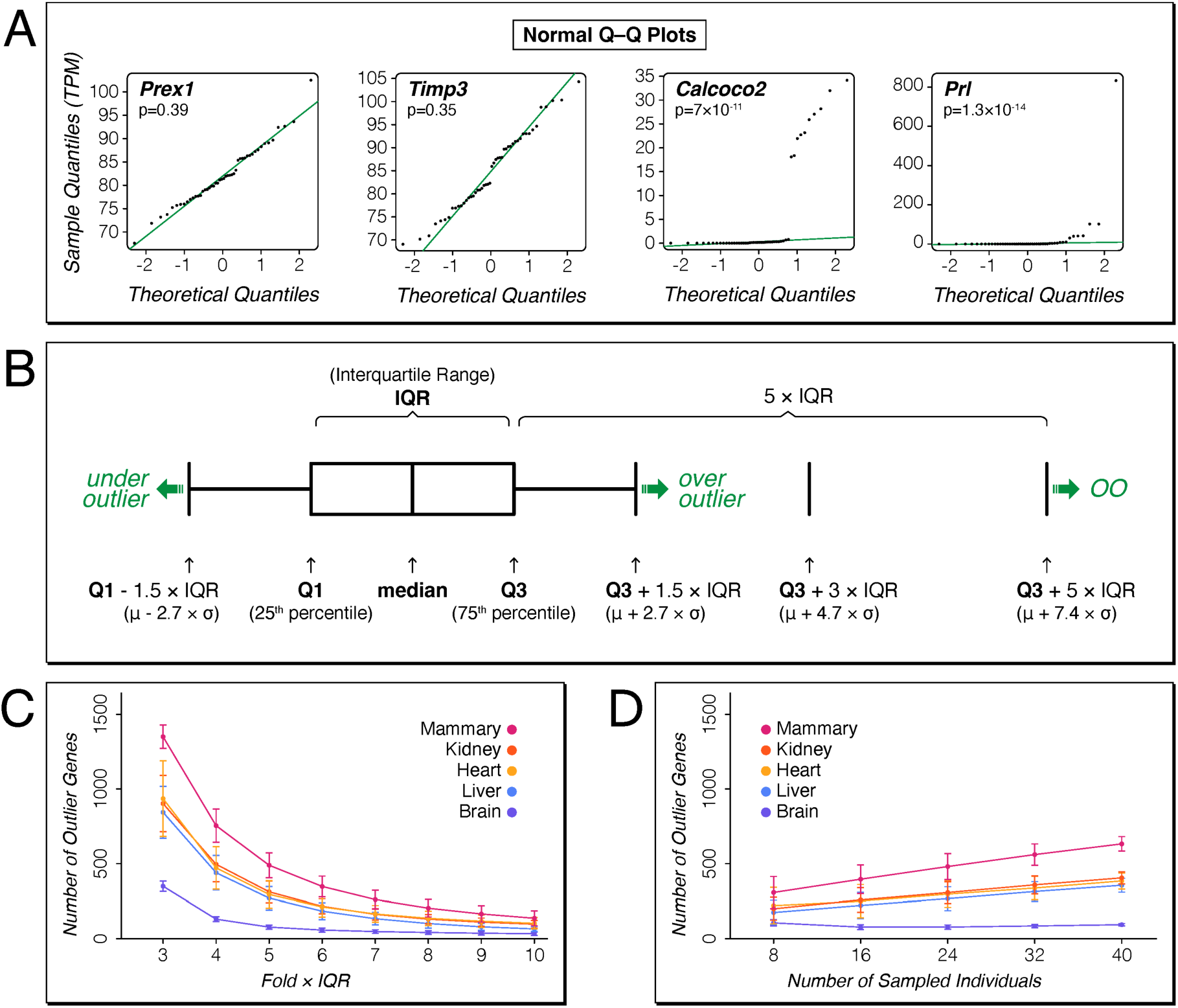
Outlier detection and analysis procedures. (A) expression data of brains from 48 mouse individuals from an outbred stock (DOM) for four representative genes plotted as Q-Q plots, with the theoretical quantiles on the X-axis and the sample quantiles on the Y-axis. P-values are calculated with the Shapiro-Wilk normality test. There was no significant deviation from the normal distribution for two of the genes (left panels), but a highly significant deviation for the other two genes (right panels). (B) Scheme for using multiples of the interquartile range (IQR) as cutoffs to identify outliers. Under outliers and over outliers are usually called at 1.5 × IQR below or above the first (Q1) or third (Q3) quantile, respectively, which corresponds to 2.7 standard deviations from the mean for normal distribution. We focus our study on over outlier expression, based on Q3 + 5 × IQR, which corresponds to 7.4 standard deviations above the mean for normal distribution. Expression values above this point are denoted as over outliers (OO) throughout. (C) Number of putative genes with at least one OO at different fold × IQR cutoff levels; averages with standard deviations based on 1,000 re-sampling rounds of 24 individuals each from the mouse dataset. (D) Number of putative genes with at least one OO at the Q3 + 5 × IQR level; averages with standard deviations based on 1,000 re-sampling rounds of different numbers of individuals each from the mouse dataset.

Here, we are particularly interested in these outliers. There are multiple possibilities to generate statistical cutoffs for such outlier detection. One could simply use a cutoff based on multiples of the standard deviation of the distribution. However, this would imply the assumption of a normal distribution of the data, which is not given in the case of multiple outliers, as shown in Figure 1A. Instead, we use the interquartile ranges (IQR) around the median of the expression values, which are less affected by skewness and extreme values and reflect the dispersion of the data. In this statistic, Tukey’s fences method (14) identifies outliers as data falling below Q1 - k × IQR or above Q3 + k × IQR, where Q1 and Q3 are 1st and 3rd quartiles, respectively (Figure 1B). For reference, when compared to a normal data distribution, k = 1.5 would correspond to a 1% cutoff, or 2.7 standard-deviation above the mean (P-value approx. 0.069). This is usually considered already as very stringent, but not sufficient when very many comparisons are done, as it is typically the case for transcriptome data with thousands of tested genes. It is therefore common to use a k = 3, which would correspond to 4.7 standard deviations above the mean (P-value approx. 2.6×10^-6^), which would satisfy even a stringent Bonferroni correction for multiple testing.

To assess the effect of different k-values on the cutoff for calling outlier values, we used resampling of the mouse dataset of 48 DOM individuals to assess which average percentage of genes would qualify as extreme outliers for different k-values in repeated sub-samples of 24 mice from the total dataset. The distribution showed that at k = 3, about 3% - 10% of all genes (∼350-1350 genes) exhibit extreme outlier expression above the overall expression in at least one individual; these results were similar across different tissues (Figure 1C). The numbers of extreme outliers continuously declined with increasing k, without a clear cutoff being identifiable (Figure 1C). For the further analysis we chose a k = 5 (i.e., Q3 + 5 × IQR) as a threshold for very conservatively defining extreme over-expression. For comparison, this cutoff corresponds to 7.4 standard deviations above the mean in a normal distribution (P-value approx. 1.4×10^-13^).

In the following, we denote extreme over-expression in a given individual as OO (compare Figure 1B) and genes harboring an OO in any individual are designated as “outlier genes”.

Given that OOs are mostly found in only one or few individuals in each dataset, the number of outlier genes is expected to depend on the total number of individuals sampled. We assessed this by random down-sampling from the larger set of 48 DOM individuals. As expected, numbers of genes with extreme expression go down with sample size, but even with only 8 individuals sampled, one can still detect about half of the genes (Figure 1D). For the following analysis, we focus mostly on datasets with large numbers of individuals (N > 40) for the primary analysis, but include also comparisons with smaller numbers of individuals.

### Distribution of OOs among individuals

The genes with OO expression were systematically determined for each individual for the whole transcriptome datasets according to the above criteria (i.e., the Q3 + 5 × IQR cutoff).

The results are summarized in Table 1 (the detailed data lists are provided for the mouse DOM and the human data in suppl. Table S2A-I and all data are included in suppl. Data S1). The majority of individuals sampled carried at least one outlier gene in at least one organ.

**Table 1:**
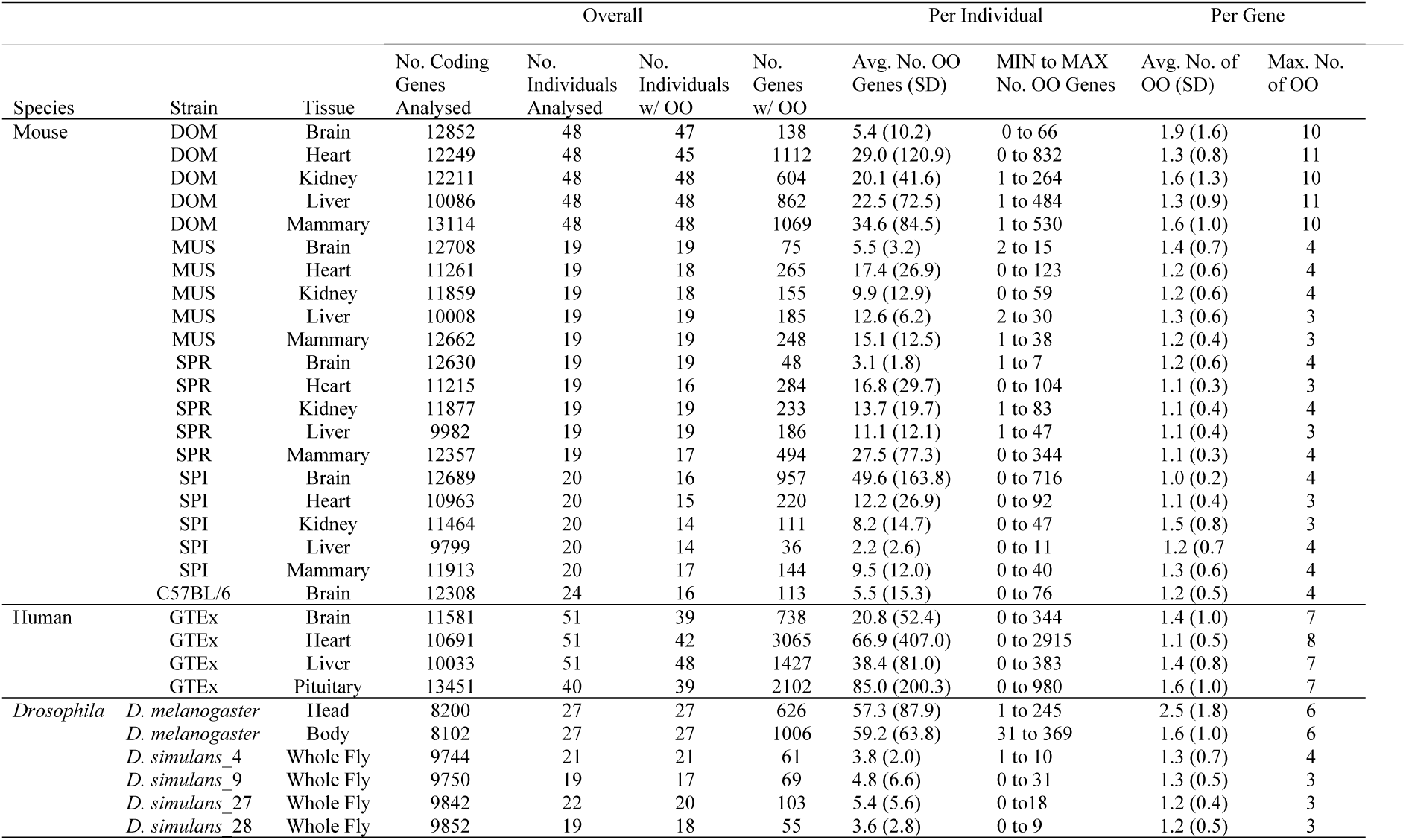
Summary statistics for the outlier calls for all species and strains.

The overall number of outlier genes per population sample ranged from 36 to 3065, with major differences between species and organs. Major differences were also found at the individual level, with some individuals harboring only few outlier genes, others very many (averages between 2.2 to 85). In contrast, at the gene level, the average number of OOs per gene is similar (averages from 1.0 to 2.5).

The comparison of individuals within any particular sample group shows extreme differences of outlier gene numbers between the individuals. This is evident from the large SD around the average and the large spread between minimum and maximum number of outlier genes per individual (Table 1). In Figure 2, we visualize this for the mouse DOM data and human data from overlapping organ sets (data are listed in suppl. Table S2). For example, individual “18” in the mouse data harbors already 832 out of the 1,112 outlier genes for the heart (= 75%), but has average numbers of outlier genes for the four other organs. Similarly, individual “27” in humans harbors 2,915 out of 3,065 of all outlier genes found in the heart (=95%), but it has only 10 and 22 outlier genes in the brain and the liver. In the following, we denote individuals with particularly many outlier genes as “outlier individuals”.

**Figure 2:**
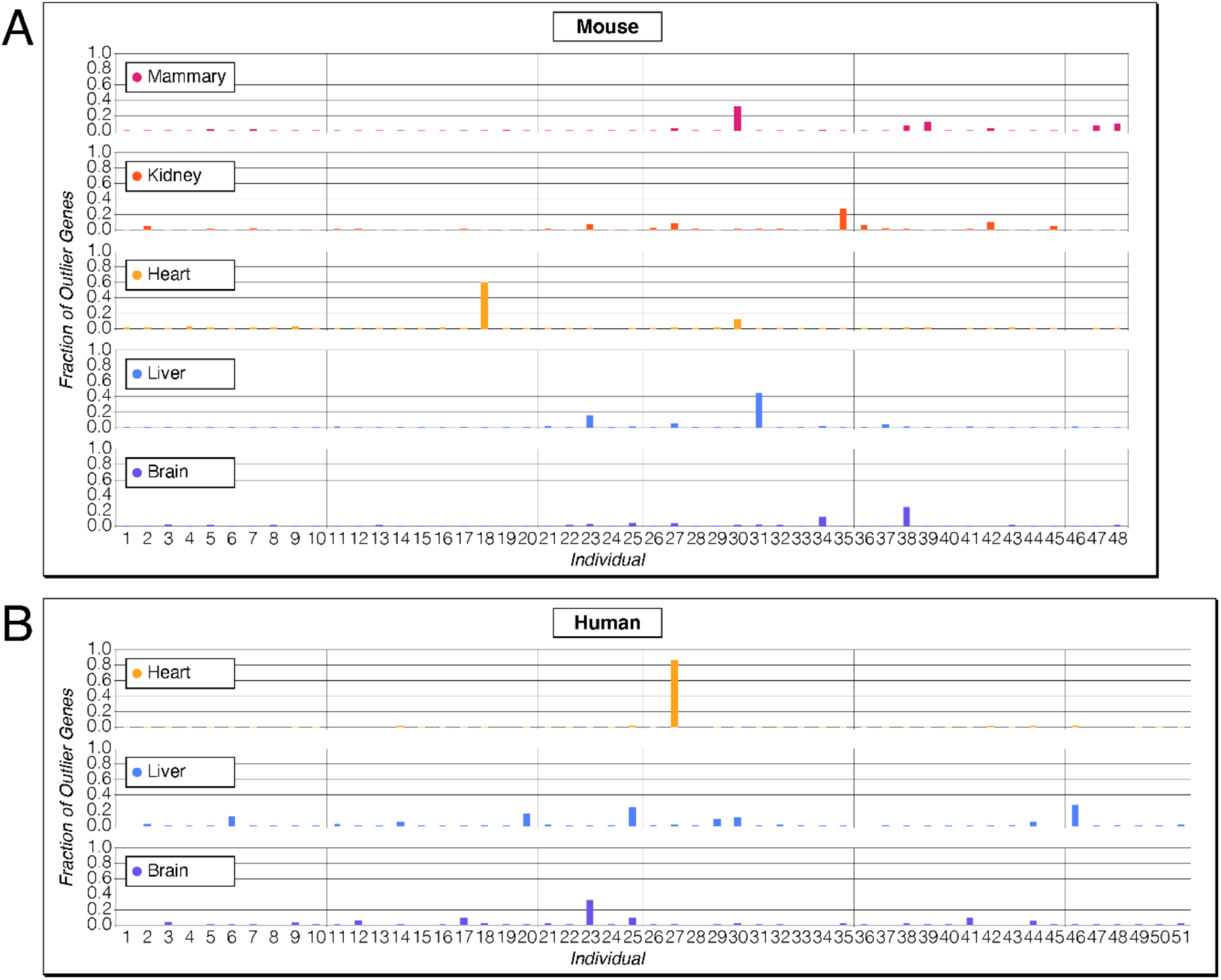
Depiction of the fraction of outlier genes per individual for organs and tissues. The upper panel is for the mouse dataset, the lower panel for the human dataset. The x-axis represents the different individuals, the y-axis represents the per individual fraction of outlier genes for the given tissue. The highest bars represent “outlier individuals” that show a particularly large number of outlier genes. The numbering of the individuals corresponds to the numbering provided in suppl. Table S2.

### Distribution of outlier genes among tissues and species

A subset of genes shows OO expression in more than one organ. In the mouse DOM data, these are 18% (565 out of 3063; suppl. Table 3A) and in the human data 22% (908 out of 4185; (suppl. Table S3H; note that only the three organs with the full set of individuals were included in these comparisons). This shared OO expression between organs could be due to a genetic polymorphism. If this were the case, it should occur in the same individuals for two or more organs, but we find that this is not the case for the majority of them. In suppl. Table S3B (DOM mouse) and suppl. Table S3I (human) we list all outlier genes that show OOs in more than one organ and identify those genes that could be due to genetic polymorphisms since they show outliers in different organs of the same individuals. In the mouse data, we identify 61 out of 565 such genes and in the human data 32 out of 908 (all highlighted in yellow in suppl. Tables S3B and S3I). This implies for most OOs that it is unlikely that they are triggered by genetic polymorphisms.

A subset of outlier genes shows OO expression in more than one mouse taxon. Between 7% - 17% of outlier genes in a given organ were found in two or more taxa (suppl. Tables S3C-S3G). When comparing all mouse OO genes found in any organ and taxon with the list of all OO genes found in humans in any of the three common organs, we detected 929 gene pairs of orthologs between mice and humans (suppl. Table S3J).

### Technical replication

Given the somewhat scattered occurrence of most OOs, one can ask whether this might be a technical artefact of the library preparation or sequencing reaction. The methodology is generally assumed to be sufficiently robust in particular compared to the microarray technology (8). Accordingly, there is a consensus that technical replicates are not required in RNA-Seq experiments. In fact, by counting multiple independent sequencing reactions in a given run is already considered to constitute an internal technical replication, which may only generate reproducibility problems when the total number of counts becomes too low, for example in single-cell sequencing experiments. Similar considerations apply to the library preparation. While this could be biased towards favoring certain sequence types, it should not generate OO patterns for identical RNAs in different individuals.

To verify these general assumptions, we generated for five individuals two separate sequencing libraries from the same brain RNAs (from the pedigree animals described below), but using different kits and sequenced the libraries on different instruments (Illumina NovaSeq and NextSeq). A total of 17 outlier genes were found in these five animals, in different combinations. Each was faithfully detected as OO in both replicates, even those with relatively low TPM values overall (suppl. Table S4). Hence, although this was only a small replication experiment, it generated no indication that OO calls could be artefacts of a given library preparation or sequencing instrument. In fact, the results from the repeated finding of the same outlier genes in different organs and species (see above), as well as the family analysis and the co-expression modules described below support further the notion that the expression measurements via RNA-Seq are a faithful method to trace OOs.

### OO expression in an inbred strain

Given that the data above suggest that OO expression could be a spontaneous phenomenon, independent of genetic polymorphisms, we asked whether we would observe OOs also in transcriptome data from an inbred mouse strain. To assess this, we obtained brain transcriptome data from 24 individuals of the isogenic lab strain C57BL/6 (suppl. Data S1E). 16 of these showed an OO expression, involving 113 genes (Table 1). Most interestingly, among the C57BL/6 mice, we also found an outlier individual with 76 outlier genes.

### Inheritance patterns of OOs in pedigree data

To further test whether genetic polymorphisms, or inherited epigenetic signals could be a key driver of OO expression, we set up five three-generation families of an outbred stock of *M. m. domesticus* originally collected from Germany. The breeding schemes are shown on the top of Figure 3 and in suppl. Table S5A). Brain, kidney and liver transcriptomes were generated for each individual and analyzed as described above. The distribution of the OO patterns across the family structures are shown in Figure 3 for the brain data, the data for all three organs are provided in suppl. Tables S5B - S5D. We identified a total of 123 (brain), 178 (kidney) and 659 (liver) outlier genes in the total dataset of 50 individuals, which is in the range of the number of outlier genes found for the first set of DOM individuals shown in Table 1.

**Figure 3:**
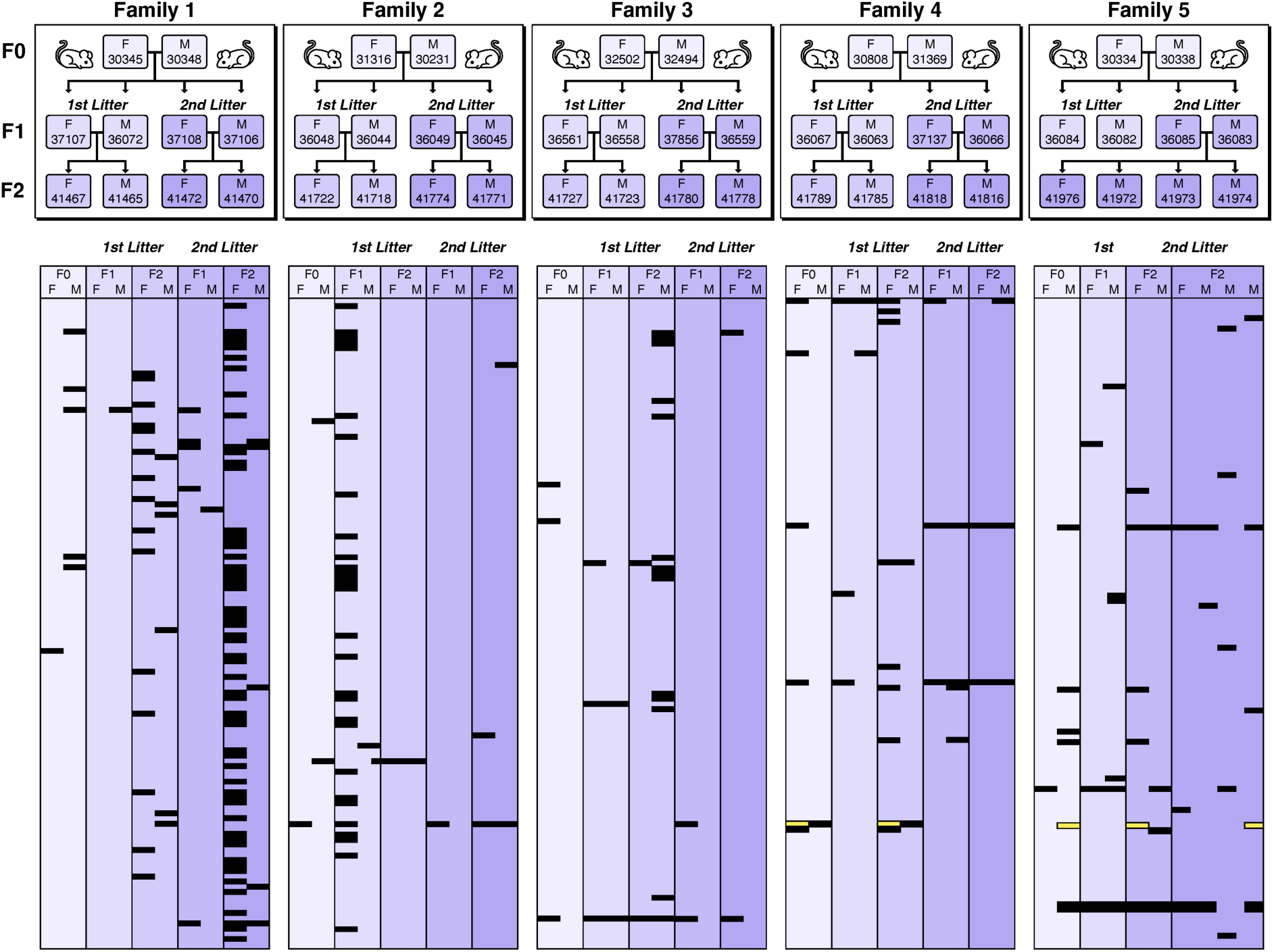
Depiction of the five mouse family pedigrees and OO patterns for brain. The upper panels show the pedigrees, whereby families 1 - 4 have the same scheme (see also overview in suppl. Table S5A). In family 5, the F1 of the first litter had no offspring. To compensate for this, we used four offspring of the F1 of the second litter. The lower panels depict rows for each gene that showed an OO in at least one individual. The OOs are highlighted in black, yellow indicate under outliers (UO - note that genes that showed only UOs are not included). The parental columns are always first in each block, followed by two F1 and two F2 individuals, for each litter. TPM values are available in suppl. Tables S5B-S5D.

Expression levels of genes should normally add up between the two alleles when they are under cis-regulation control (15). This will show up as an additive (co-dominant) inheritance. In case of trans-regulation, it is a question of the presence or absence of the trans-regulator, which will also show up as dominant inheritance pattern. In both cases, one does not expect a recessive pattern. Based on these considerations, we checked whether the distribution of OOs in the families were compatible with a Mendelian dominant (or co-dominant) inheritance. We classified all genes into *Mendelian (M)*, when the inheritance patterns of the OOs were compatible with Mendelian inheritance in at least one family and as *Spontaneous (S)*, when OO turned up in offspring, but not in their parents. We used also two further classifications where inheritance assignment was not possible (*P* for *Parental*, i.e., only occurring in the F0 and *Q* for *Questionable*, i.e., not clearly resolvable). The classifications are based on an automated algorithm and were combined with a manual analysis according to the criteria detailed in the Methods section.

In this gene-by-gene analysis, we found that the majority of OOs could be classified as having spontaneously arisen (brain 101 *S* vs. 10 *M*, kidney 87 *S* vs. 15 *M*, liver 158 *S* vs 17 *M*) (suppl. Table S5B-D). For the *M* genes that were expressed in more than one organ, we found that the OO pattern was usually found in the same individuals of the other organ (suppl. Table S5E), confirming the notion of a genetic polymorphism driving the pattern.

An interesting *Q* classified gene in the brain data is *Glyoxylase 1* (*Glo1)*. This is a gene known to have high copy-number variability in the DOM populations and its expression level is known to influence anxiety patterns (16). It shows variable expression levels among the individuals, of which only some may be inherited. It is also the only gene that shows both, OO as well as under-expression (under outlier, UO) (Figure 3 and suppl. Table S5B). This suggests that new copy number variants are frequently generated in these family pedigrees. Among the other genes classified as *Q* in the different organs, we found only *Rnase2a* as a gene that has previously been identified as being copy number variable in the dataset of (17), while the others are single-copy genes.

There are also at least four individuals in the family set which can be classified as “outlier individuals”, based on harboring particularly many outlier genes. Figure 3 shows this for two individuals (individual 41472 in family 1, F2, second litter and individual 36048 in Family 2, F1, first litter). In both cases, the parents did not have large numbers of OOs, suggesting that the status as “outlier individual” has arisen spontaneously in the respective offspring. In the liver data, there are two parental F0 animals in family 1 and family 2 (individuals 30345 and 31316 - suppl. Table S5D) that constitute such “outlier individuals”. Both did not inherit this status to the F1 in the respective families, which is also compatible with the assumption that the status as “outlier individual” is a spontaneous one.

### Correlated OO expression

During the analysis of the data, we noted for some gene sets a correlated OO expression in their respective tissue. To assess this systematically, we focused on genes in which at least three individuals sharing the OO expression for two or more genes and no other individual showed an OO expression for the respective genes. This is a rather strict requirement, but even under this condition, we find many such correlated expression groups. As an example, we show in Figure 4 a subset of the groups and distributions between individuals for the mouse organ data. This represents the cases where at least four individuals share an OO expression, the extended version with at least three individuals sharing an OO expression is provided in suppl. Table S6A.

**Figure 4:**
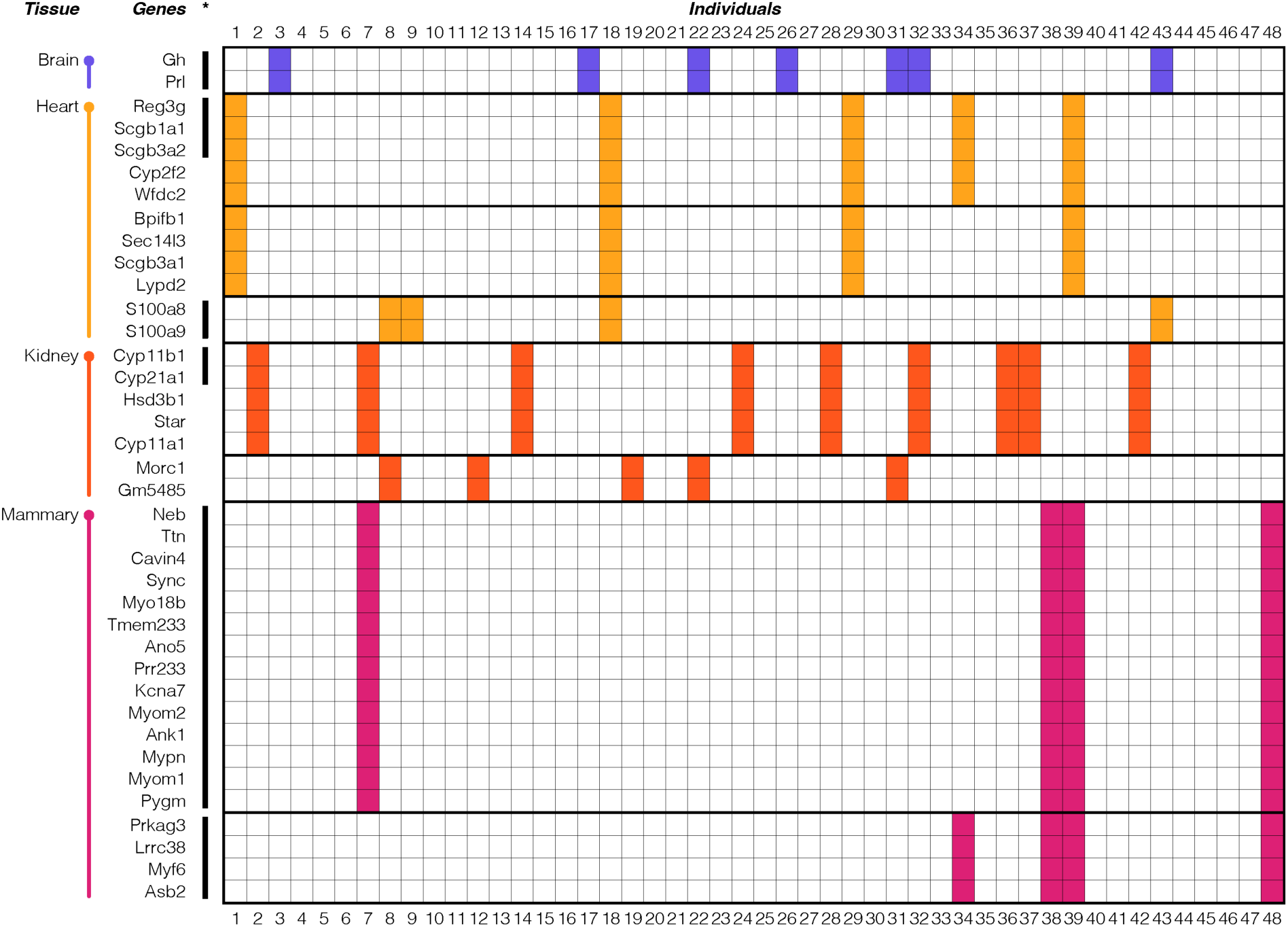
Correlated OO expression in gene groups in mouse organs. Genes are shown for which the pairwise comparisons revealed at least four individuals with OOs and no individual gene with OO expression in another mouse (an extended version of this figure with a minimum of three individuals sharing OO expression is provided in suppl. Table S6A). OO expression is marked by colored shading. Note that liver shows no such gene pairs or gene groups, and thus is not included. Black lines to the left indicate co-expression groups that are still visible when the outlier individuals are removed from the respective comparisons.

OOs in Figure 4 are marked by colored shading (colors according to tissues), outlier gene groups by boxes. In the heart, there is the gene pair S100 calcium binding proteins A8 and A9 (*S100a8* / *S100a9*), which is located in the same chromosome region, but on different strands. The genes code for highly studied Ca-binding proteins, with a role in inflammation and cancer (18). They act usually as heterodimers, i.e., a correlated expression is expected for them.

Heart includes further a group of five outlier genes, of which two are members of a family of Secretoglobins on different chromosomes (*Scgb3a2* and *Scgb1a1*), one a member of the Cytochrome P450 family of proteins (*Cyp2f2*), one a member of the Regenerating islet-derived gene family (*Reg3g*) and the WAP four-disulfide core domain 2 gene (*Wfdc2*). In kidney, there is also a group of five outlier genes, three of them part of the Cytochrome P450 family (*Cyp21a1*, *Cyp11b1* and *Cyp11a1*, all encoded on different chromosomes), in conjunction with a Hydroxy-delta-5-steroid dehydrogenase gene family member (*Hsd3b1*) and the Steroidogenic acute regulatory protein (*Star*). To assess whether these groups may have joint functions, we used GO analysis via PANTHER (19), but without significant results. On the other hand, the top block of 14 co-regulated genes in mammary, are enriched for genes involved in myofibril assembly including Myomesin 1 and 2 (*Myom1*, *Myom2*), Myopalladin (*Mypn*), Caveolae-associated protein 4 (*Cavin4*), Titin (*Ttn*) and Nebulin (*Neb*).

Note that the group modules show frequently partial sharing of individuals with other group modules (e.g., visible in heart and mammary in Figure 4, and more extensively visible in the extended supplementary Tables with the full data for mice, humans and *Drosophila* - suppl. Tables S6A-S6C).

We asked whether these co-regulated outlier gene groups reflect modules that are also co-regulated when they include no individuals with OO expression. To assess this systematically, we removed the individuals with OO from the data and asked whether the remaining gene expression levels between the individuals was significantly correlated (Kendalĺs tau for every combination of gene pairs, P<0.05 with Bonferroni multiple test correction). It is indeed possible to find correlated gene groups in this way which overlap at least partially with the gene modules identified via over-expression analysis. These are marked by black lines in Figure 4 and green boxes in suppl. Tables S6A-S6C. Interestingly, the gene group of 14 genes in mammary that may be involved in myofibril assembly shows also up as co-regulated group when OOs are removed.

Other correlation groups are only partly visible when the OOs are removed, e.g., the two groups of five genes discussed above for heart and kidney (Figure 4), or even absent (compare also extended data in suppl. Tables S6A-C). In the *Drosophila* data there are very large and complex groups of outlier genes with sharing between individuals, especially in the head, but very few of them show co-regulation when the OOs are removed (suppl. Table S6C).

We conclude from this analysis that outlier genes that are co-expressed in multiple individuals identify to some degree co-expression modules, which may, or may not be detected by other co-expression analyses as well.

### Outlier in single cell data

Gene expression is generally known to be pulsatile, whereby only a subset of cells in a tissue show high expression at a given time (20, 21). We have therefore also analyzed single-cell data to trace the phenomenon of over-expression in some individuals. However, because one needs the comparison between multiple individuals, there are still only few datasets that can be used for this purpose. We have chosen data from an Alzheimer study, which has investigated two human brain regions (DFLPC and MTG) from many individuals using single-nucleus transcriptomics (22). We include a total of 53 individuals and up to 15 cell types that were consistently identified between them (suppl Table S1F), each covered by at least 100 cells per individual. Given that the sequencing depth at each single cell is relatively low, we used first the mean across all cells for the given cell type in a given individual for the overall analysis.

The overall analysis is comparable to the data from the organs described above. We found between 0 to 68 OOs per individual (average 5.8, SD = 12.5; 0 to 26 outlier genes, average 3.4, SD = 5.9) and 2 to 48 OOs per cell type (1 to 40 outlier genes) (suppl. Table S7A). The numbers between the individuals differ substantially and the two brain regions show different numbers for the same individuals (Figure 5A). There are also major differences in OO genes between the cell types, whereby the numbers are similar for those cell types that occurred in both brain regions (Figure 5B).

**Figure 5:**
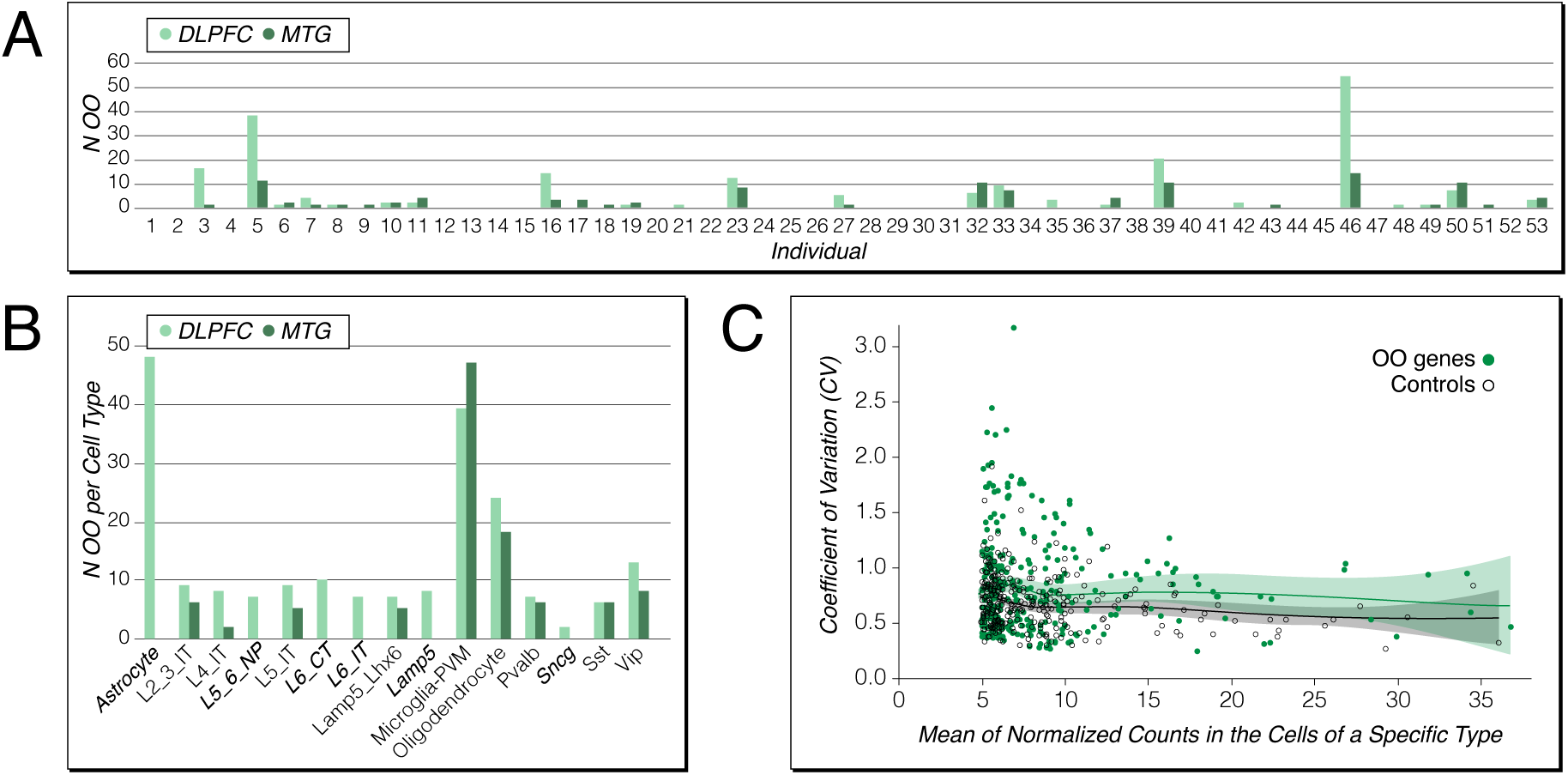
Numbers of OOs in individuals and cell types and spread of variances of OO genes in single nuclei expression data. (A) Cumulative numbers of OOs for the cell types from each of the two sampled tissues, for all individuals. (B) Number of OOs per cell type. Note that MTG includes only a subset of the cell types of DLPFC (suppl. Table S7A), the cell types specific for DFLPC are written in italics. (C) Spread of variances of OO genes compared to control genes. Every OO gene is matched with a control gene from the same cells, which is expressed at a similar level, but without much variance between the individuals. The two distributions are significantly different (P = 1.2✕10^-6^, Wilcoxon signed rank test).

However, the power of the single-cell data lies in asking whether the OO status across the mean of the transcript counts is generated more or less by all cells, or by a subset of particularly highly expressing cells. To test for this, we focused on all 145 genes that showed an OO in at least one individual and in one or more cell types. We asked whether the variance in expression in the individual cells for a given cell type would exceed the variance of a control gene in the same set of cells. Control genes were chosen such that they had a similar overall expression level as the OO gene, but a variance within the Q1-Q3 range across individuals. Given that a number of the 145 genes were expressed as OO in more than one cell type, we could do a total of 307 such comparisons (suppl. Table S7B). We found that the overall variances (measured as coefficient of variation = CV) was significantly higher across the OO genes than across the control genes (P = 1.2×10^-6^, Wilcoxon signed rank test), but with substantial overlaps (Figure 5C). In fact, only 176 comparisons have a higher CV, 131 a lower one (suppl. Table S7B).

### Prolactin and growth hormone

Among the outlier genes shared between mice and humans are the hormone genes, prolactin (*Prl* / PRL in humans) and growth hormone (*Gh* / GH1 in humans). They belong to the co-regulated gene pairs described above, with the same individuals displaying the OO expression status for both genes in the outbred and inbred mice, as well as in humans (Figure 6). In humans, both genes are also expressed in the heart and they show also OOs for this tissue, but in different individuals compared to the brain (Figure 6C,D). *Gh* and *Prl* expression is also significantly correlated when the OO individuals are removed, both in the mouse brain (Figure 4, top) as well as in the human brain and heart (suppl. Table S6B).

**Figure 6:**
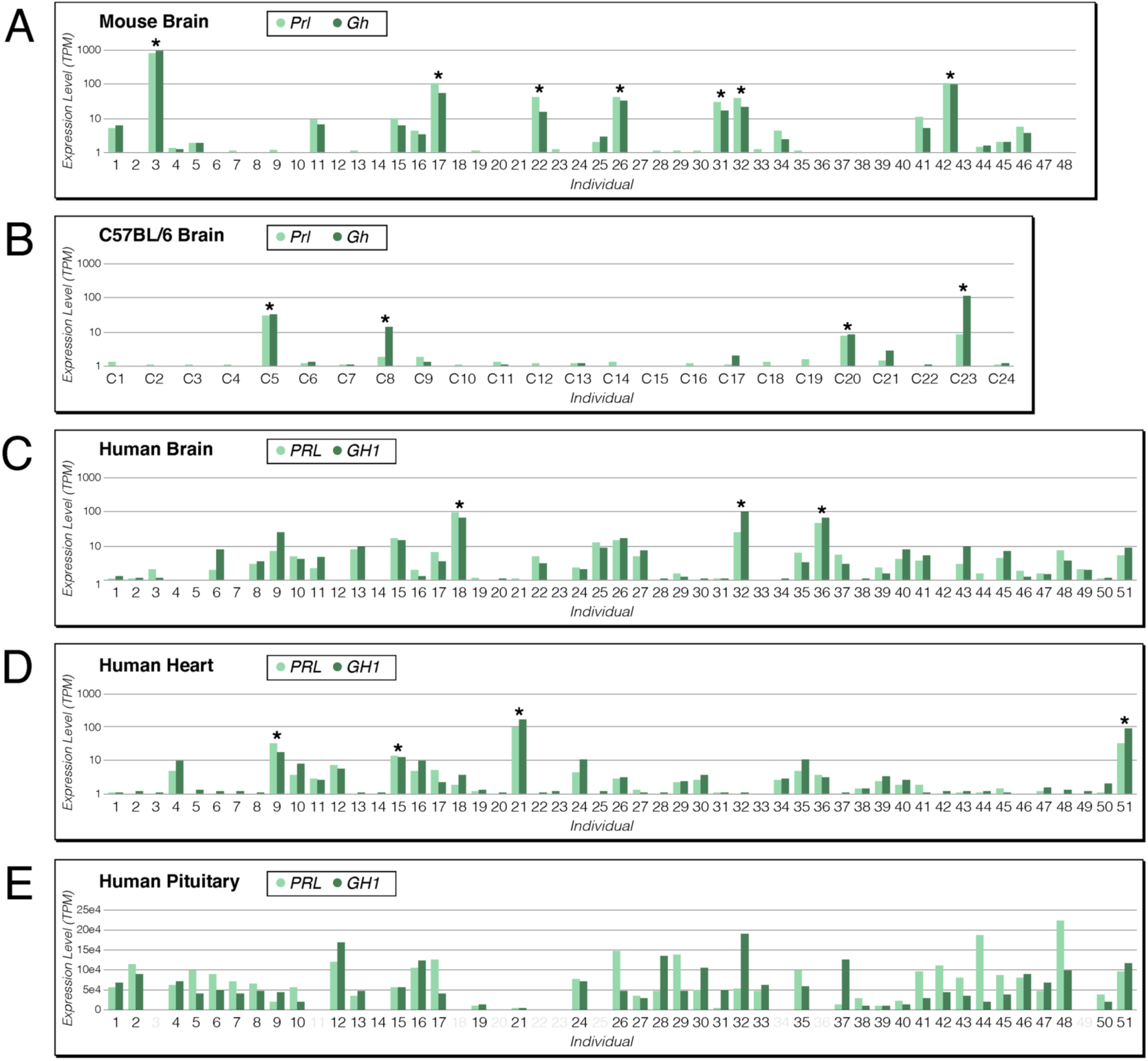
Expression levels of prolactin (light green) and growth hormone (dark green) transcripts between individuals. Data include mice (A and B) and humans (C to E). The Y-axis for (A-D) is on a log scale of “TPM + 1”. (A) DOM wild mouse brain, (B) isogenic mouse strain C57BL/6 brain, (C) human brain, (D) human heart, and (E) human pituitary gland. OOs meeting the Q3 + IQR * 5 threshold are marked with stars. Note that the panels for humans relate to the same individuals (51 for brain and heart, 40 for pituitary gland), i.e., OOs occur in different individuals for brain and heart.

Hence, the OO expression of this gene pair reflects an inherent co-expression of these genes in the respective tissues.

However, the expression of these hormone genes in the general brain and heart tissues is relatively low and of unknown function. The main expression of the genes occurs in the pituitary gland, but in different cell types, the somatotroph and lactotroph cells, respectively. We have therefore also examined the GTEx data for the human pituitary gland from the subset of individuals that overlap with the main human GTEx dataset described above. The analysis is included in Table 1.

While the human pituitary gland shows generally a large number of outlier genes, PRL and GH1 are not among them. However, there are substantial expression differences between the individuals (Figure 6E), implying a large IQR spread, such that some highly expressing individuals (e.g., individuals 44 and 48 for PRL, or individual 32 for GH1) fall below the chosen cutoff. Interestingly, there is also no correlation between PRL and GH1 expression levels in the pituitary gland (Figure 6E), as is to be expected given that they are expressed in different cell types of the gland.

Hence, the expression characteristics of the two hormone genes in the secondary tissues (brain and heart) are different from the primary tissue (pituitary gland). This implies that the overexpression in the secondary tissues is not simply the consequence of using the same promotor signals as in the primary tissue. Of note, *Prl* and *Gh* showed also up as OO in the mouse family brain data (suppl. Table 5B), both in the same four individuals, *Gh* in one additional individual. In three out of the five cases, there was no parental individual from which the expression could have been inherited, i.e., these would be classified as *S*. However, in Family 5, litter 2, an inheritance could not be excluded. Hence both genes were conservatively classified as *Q* (suppl. Table S5B), although *S* (spontaneous activation), as is suggested by the OO pattern in the inbred mouse strain (Figure 6B), is also compatible.

### Outlier in hormone level measurements

Prolactin and growth hormone are often measured in large human cohorts. A recurrent observation in these studies is the identification of individuals with exceptionally high hormone levels (23, 24). While high hormone levels might be caused by a medical condition (25), we analyzed the data under the aspect of OO expression patterns reflected at the protein level.

All the above data are from RNA measurements, which represent only a single point in the lifetime of the individual. It is therefore not possible to determine whether outlier expression status is a short-term phenomenon (in the order of days or weeks), or whether it is stable for a given individual over the lifetime. This is different for hormone measures, which can be done repeatedly for the same individuals.

We evaluated plasma concentrations of prolactin and growth hormone measured as part of case-control studies for breast cancer nested within two large prospective cohort studies, the Nurses’ Health Studies (NHS/NHSII). Details of assay measurements and quality control are available in (26, 27). For outlier status we used the Q3 + 3 × IQR cutoff, since the multiple testing problem is not an issue here (only two hormones tested, in contrast to thousands of genes in the RNA analysis above) and since the Q3 + 5 × IQR cutoff appeared to be stringent for the RNA data (see above).

In one study (NHS), data were obtained from 310 females, with two measurements of prolactin approximately 9-15 years apart (first blood sample collected from 1989-1990 and second from 2000-2003). The overall correlation between the two measurements was moderate, but significant (0.27; Kendalĺs tau; p < 0.0001). Four individuals showed outlier levels, three in the first measurement, one in the second, but none showed outlier levels at both times (Figure 7A).

**Figure 7:**
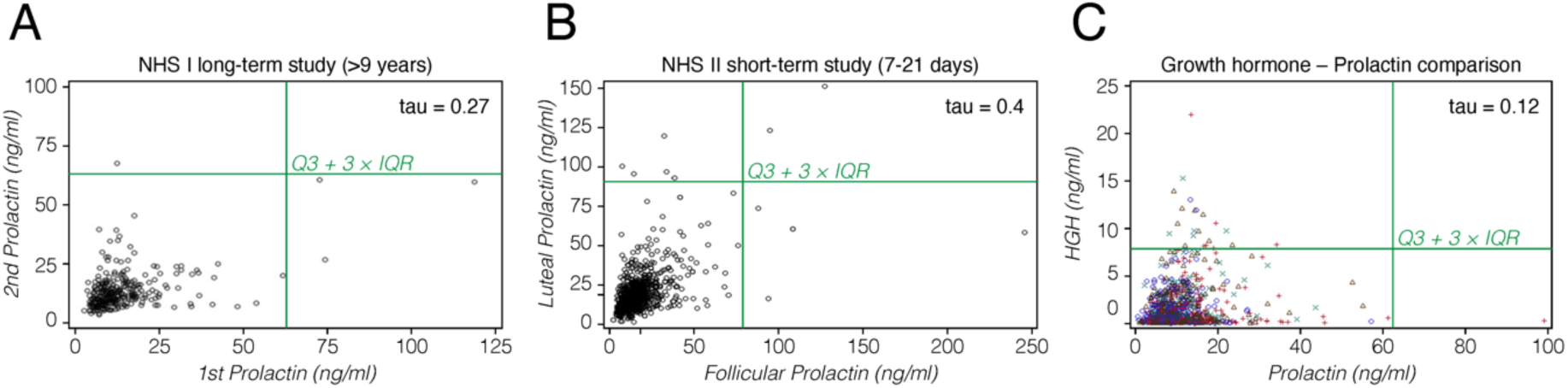
Hormone expression level comparisons in humans. Results are shown for prolactin measurements at two different times (A, B), and a comparison between prolactin and growth hormone in the same individuals (C). (A) Data from 310 females for which measurements were taken between 9.5 to 15.25 years apart. (B) Data from 974 females for which measurements were taken during one menstrual cycle, the first in the follicular phase, the second in the luteal phase. (C) Data from 1,200 females, taken at different ages for prolactin and growth hormone (HGH). Different symbols represent different ages that are not further considered here.

In the second study (NHSII), 974 females were tested during a menstrual cycle, with first measurement in the early follicular phase (about 5 days after onset of menses) and the second in the luteal phase, i.e., about 7-9 days before the anticipated start of the next period; the samples were collected within the same cycle (within the same month). This tests for short term stability of outlier levels. The correlation within person for these values was higher than for the long-term study (0.4; Kendalĺs tau; p < 0.0001). Eleven individuals with outlier levels were detected, four in the follicular phase only, five in the luteal phase only, and two in both phases. Individuals with outlier levels in one phase generally showed elevated hormone levels at the other stage, although below the cutoff (Figure 7B).

In NHS/NHSII, a subset of women with prolactin measures also had growth hormone measures on the same blood sample. In the NHS data, 22 individuals with outlier levels were detected for growth hormone among 1,200 females in NHS, but only one individual with outlier level for prolactin, and this individual did not show an elevated growth hormone level (Figure 7C). Nonetheless, there was still a low, but significant overall correlation across all data (0.12; Kendalĺs tau; P < 0.0001).

To put this into context with the RNA data of pituitary glands, we analyzed all 283 individuals from the GTEx study. In this larger group and with a Q3 + 3 × IQR cutoff, three individuals had PRL with an OO, but still none had one for GH1. The overall correlation across all data was similar to the protein data (0.12; Kendall’s tau; P = 0.0021).

These observations show that the strongly correlated patterns of OO expression seen for the two hormones in secondary tissues are not reflected in the circulating hormone levels. On the other hand, the RNA expression patterns from the primary tissue (the pituitary gland), are well compatible with the protein data.

## Discussion

The occurrence of outliers in expression values is a well-known phenomenon in transcriptome data. However, this has so far been treated as a noise component of the data and it was corrected by filtering techniques, such as removal of certain samples, or transformation and/or modelling of the data spread to reduce the relative impact of outliers on the overall analysis.

Here we have studied RNA expression across multiple species, including mice, humans, and *Drosophila* to better understand the occurrence and patterns of outlier expression. We use an interquartile range (IQR) statistic without data transformation to identify extreme over-outlier (OO) in the datasets that are far beyond expected statistical fluctuations. We show that the detection of such outliers is reproducible between different library preparation protocols and different sequencing machines.

We find that OOs are distributed unevenly across individuals, with hundreds to thousands of genes with an OO in one or a few individuals in a dataset. Some individuals harbor particularly many outlier genes in a given organ but not in another organ from the same individual. Most of these outliers are not driven by genetic polymorphisms, as seen in both inbred mouse strains and a three-generation family study in mice.

OOs can occur in co-regulated modules, in the sense that they are co-expressed in multiple individuals. Several of these modules reflect also a co-expression of the same genes when they are expressed in a normal range. However, some modules show up only in the outlier gene expression, but not in the normal expression. This effect is particularly evident in the *Drosophila* data.

### Prolactin and growth hormone gene expression

Among the co-expressed genes with frequent OO patterns in mice and humans are the hormone genes prolactin and growth hormone. While the highest expression of these hormones is in the pituitary gland, they are also expressed in multiple other tissues, including the brain cortex and the heart, but at much lower overall levels. In the mouse brain preparations, the pituitary gland is included in the whole brain. But given that it is so small, the RNA derived from it would be highly diluted by the RNA from the other brain areas.

Hence, we assume that the expression that we can detect in mouse is mostly from the general brain tissue. For the humans, we could separately analyse the pituitary gland and it harbors indeed many genes with OO patterns. PRL and GH1 expression is approx. 10,000 times higher in this tissue than in any other tissue. However, both of these genes did not show an OO pattern in any of the 40 analysed individuals, at least for our strict cutoff criteria. Still, there is major variation in expression of these genes between individuals, hence the general effect that leads to OO patterns could still be active. Interestingly, there is only a weak correlation in expression levels of PRL and GH1 in the pituitary gland, in contrast to a strong correlation in the brain cortex and the heart. This suggests that the regulatory input into these genes is different between the peripheral tissues and the main tissue of expression. On the other hand, the expression correlation in secondary tissues is conserved between mice and humans, implying that it is not a chance effect. These hormones have rather pleiotropic functions across the body and in various physiological processes (28) and additional functions in tissues where they are co-expressed at low levels seem possible.

### Longitudinal analysis

The longitudinal analysis of the prolactin hormone data in humans suggests that the OO expression may be somewhat stable within a matter of days or weeks. However, currently this can only be a first attempt to approach this question. Given that the RNA expression in the pituitary gland did actually not show up prolactin to be an OO gene, we cannot even be certain that the large variation of the prolactin expression is actually based on the same principles as the OO expression. High prolactin expression may also be caused by adenomas (25).

Ideally, longitudinal studies should be done based on RNA-Seq experiments to screen all expressed genes in parallel. There are some first developments to establish such a non-invasive sampling for RNA expression (29). Such approaches will allow to set up dedicated studies to assess how long a spontaneous OO expression lasts.

### Expression modules

Not only the prolactin growth hormone gene pair, but also several other sets of OO genes are co-expressed in multiple individuals, which serves actually as additional evidence that this is not a simple noise effect. Some of these co-expression groups would be considered as gene modules in standard correlation analyses, since they are also co-expressed when the OO individuals are removed. Interestingly, most modules of OO genes are broken up between the individuals, i.e., they occur in different, but overlapping combinations. This could be interpreted as rather short-lived phases of the OO status, which might end at different times in different individuals.

### Spontaneous activation of high expression promotors?

One could speculate that the OO incidences in given tissues are due to accidental activation of regulatory pathways that trigger a high expression of these genes in other tissues. This would predict that OO is primarily observed in tissues that have a low expression of a gene, which has a very high expression in another tissue. This would be akin to the prolactin growth hormone genes discussed above, although their co-expression in the peripheral tissues does not really support such an interpretation. Still, we tested this interpretation across all genes.

In humans, we used the list of median expression values of all genes across all 54 GTEx tissues to ask whether OO expression is more likely in tissues where they are not maximally expressed. In the liver, there are 21 OO genes that have also normally the highest expression in liver, which reflects approximately the proportion of all genes that have highest expression in the liver (1.4% vs. 2.2%). On the other hand, heart and brain have only one OO gene each that is also expressed highest in these tissues, which is a lower fraction than expected overall (heart: 0.03% vs 0.4% and brain: 0.13% vs 0.4 %). So again, the answer seems equivocal at this stage.

### Epigenetic signatures of outlier expression

One would expect that outlier expression should become visible in epigenetic signatures of gene expression, e.g. methylation patterns, openness of chromatin or polymerase loading. The EN-TEx project (30) has generated assorted epigenome data on four individuals of the GTEx samples. We have done a preliminary analysis of these data to assess whether there is any obvious pattern. We analysed ATAC-seq or DNase-seq to assess promoter openess, DNAme array data for methylation states at promoters, POLR2A ChIP-seq for polymerase loading, and histone ChIP-seq to determine histone modification states around promoters. Across the data from nine organs, we found 13 genes that had an outlier status that could be compared to corresponding data from an individual that showed no outlier expression. However, not clear consistent effect could be detected in this limited set. Only one gene showed a much higher polymerase loading, but this was not seen for the other genes. Still, once more such epigenomic data become available at a population level scale, they may eventually reveal more about the mechanisms of how the outlier expression comes about.

### Relationship to pulsatile gene expression

It is well established that expression of a gene in a tissue does usually not occur in all cells at the same time, but only in a subset of cells with particularly high expression pulses (20, 21). This has also been shown for the expression of prolactin in pituitary tissue (31). This pulsatile gene expression averages out over time across the cells in the tissue, such that a reproducible expression is measured form the whole tissue. One could therefore speculate that OO patterns are somewhat linked to such pulsatile effects. For example, if cells that are on during a pulse fail to shut off, one would get on average a higher spontaneous expression across the tissue.

Such an effect should become visible in single-cell data. For genes that show an OO in an organ from which the cells are derived, one would expect that a subset of the cells shows much higher expression than the cells from controls where no OO is observed. Alternatively, all or most cells show an elevated expression. This can be measured as spread of variance, whereby the variance should be high when only few cells express very highly. In our analysis, we measure indeed a significantly higher variance for the OO genes, but the difference to the control genes is not very large. Hence, an effect of pulsatile expression maintenance is not evident in these data.

### Edge of chaos expression?

The modeling of gene regulatory networks has revealed that complex interactions within these networks can lead to chaotic dynamics. Gene networks are often highly nonlinear and sensitive to initial conditions, two hallmarks of chaotic systems. It has long been hypothesized that living cells are at the critical boundary between an organized and a disorganized state (32) and gene expression analysis experiments in cell cultures have supported this (33).

In fact, it has been suggested that the presence of chaos in gene regulatory networks may offer some advantages, such as enhancing the flexibility and adaptability of cellular systems in complex and changing environments. Chaotic systems can show some robustness to perturbations at a broader scale, although with unpredictability at a finer scale (34). This has led to suggestions that gene regulatory network evolve towards a state at the “edge of chaos” to confer robustness against environmental perturbations (35, 36). Most interestingly, while the whole gene regulatory network could show stability, there could be sub-motives that show chaotic behavior (37). Hence, it seems possible that the outlier gene expression patterns are a signature of an edge of chaos effect in the sense that small perturbations trigger them in some individuals, while the overall network controls them. This would predict that such over-expressions are transient, putting even more emphasis on the need to generate longitudinal data of gene expression for individuals at a population scale.

### Conclusions

The occurrence of outlier expression patterns in individuals is a recurrent phenomenon across species, tissues and cells. Most of the outliers occur as singletons with a large overlap between mouse and human genes. This may imply that most, if not all genes can potentially show sporadic overexpression in some individuals. Most of these expression effects are not driven by genetic variants, but they can reflect partly co-regulated modules. They are therefore a biological reality that could reflect the fact that gene regulatory mechanisms operate at the edge of chaos.

## Methods

### Mouse data

Mouse data includes bulk RNA-Seq from wild mouse populations, an inbred mouse strain, and wild mouse families. Many of these data were part of a parallel study on sex-biased gene expression (10). In this study we found that while outlier expression can affect the statistical analysis of sex-biased gene expression, it shows itself no specific sex-related patterns. For the present study we have therefore used both, pure sex, as well as mixed-sex samples.

For the wild mouse populations, RNA-Seq data for the five different organs (brain, heart, kidney, liver, and mammary gland) were obtained together with the project described in (10). In short, the data were collected from individuals kept under outbreeding conditions (38) from two *Mus musculus* subspecies, *M. m. domesticus* from France (DOM) and *M. m. musculus* (MUS) as well as two sister species, *M. spretus* (SPR) and *M. spicilegus* (SPI). For the DOM samples, we focused on 48 females (suppl. Table S1A) which were collected with information for their estrous cycle. However, in an independent analysis, we did not find a systematic influence of the estrous cycle stage on the transcriptome patterns for the non-gonadal organs used in the current study (Xie et al. in preparation), i.e., we do not use estrous cycle stage as a factor for the further analysis. For the other outbred animals, we used both males and females, a total of 19 individuals were used for MUS and SPR, and 20 for SPI (suppl. Table S1B). Note that the five organs used were always from the same individuals. In total, data from 63 individuals are also used in the project described in (10), and the rest from 43 individuals are only used in the present project but all the data were obtained in parallel at the same time by the same people and with the same procedure.

For the comparison with the inbred strain C57BL/6, we sequenced the brain RNA from 24 animals from the corresponding inbred stock (suppl. Table S1E).

For the wild mouse families used in the outlier inheritance analysis, five three-generation families of an outbred stock of *M. m. domesticus* from Germany were set up, and each family has ten individuals. The breeding schemes are shown on the top of Figure 3 and in suppl. Table S5A. Brain, kidney and liver RNA-Seq were generated for each individual. The steps for assigning the inheritance state for each gene are described in the readme Tab of suppl. Table S5.

### Human RNA data

Human data includes bulk RNA-Seq from the GTEx project (39, 40) and single-nucleus RNA-Seq (snRNA-Seq) from the SEA-AD project (41).

The human bulk RNA-Seq data were retrieved from the GTEx project as TPM files for each organ. To make them best comparable with the mouse data, we chose three organs that were also analyzed for the mouse, brain (cortex), heart (left ventricle), and liver. We retrieved 51 individuals from which data were available for each of these organs. In addition, we also used pituitary data from 40 of the 51 individuals (suppl. Table S1C).

The human snRNA-Seq data were retrieved from the SEA-AD project as “h5ad” format files, including the normalized read counts presented as a log transformation of pseudocounts per 10,000 reads, ln(CPTT+1), which were downloaded from CZ CELLxGENE (42, 43). Cells in two brain regions were included: middle temporal gyrus (MTG) and dorsolateral prefrontal cortex (DLPFC). The data from the individuals with race as “white”, and sequenced by the “10x 3’ v3” assay was used for further selection. We selected 53 individuals and 24 cell types (“subclass”, 15 for DLPFC and nine for MTG), and each of the 24 cell types has at least 100 cells for each of the 53 individuals. The full list of cell types and individuals included is provided in suppl. Table S1F. We used CPTT as the expression level of a gene in a cell, and mean(CPTT) as the expression level of a gene in a cell type of a brain region from an individual.

### Drosophila data

*Drosophila* data includes bulk RNA-Seq from *D. melanogaster* (12). and *D. simulans* (13). The *D. melanogaster* data includes the RNA-Seq of heads and bodies from 27 females, and the CPM files were provided by the authors. The *D. simulans* data includes the RNA-Seq of the whole flies from four populations, with 19 to 22 males of each population, and the CPM files were provided in the supporting information of the paper (suppl. Table S1D).

### RNA sequencing and data analysis

The mouse organs were carefully dissected and immediately frozen in liquid nitrogen. Total RNAs were purified using QIAGEN kits, QIAzol (Catalog no. 79306) and RNeasy 96 Universal Tissue Kit (Catalog no. 74881), and prepared using Illumina TruSeq Stranded mRNA Kit, and sequenced using Illumina NovaSeq S4 (2 x 150 bp) in Kiel Sequencing Center (for all samples except for the five technical replicates of the pedigree samples) or Illumina NextSeq 500 (2 x 150 bp) in MPI Ploen (for the five technical replicates of the pedigree samples). All procedures were performed in a standardized and parallel way to reduce experimental variance.

Raw sequencing reads were trimmed using Trimmomatic (0.38) (44) with suggested parameter settings, except we did not remove bases from the beginning of reads. Only paired-end reads left were used for following analyses. The trimmed reads were mapped to mouse genome GRCm39 (45, 46) using STAR (2.7.9a) (47). Based on the communication with the authors of STAR, the parameters were adjusted to account for the sequence divergence of the mouse taxa and the reference genome: “--outFilterMismatchNmax 30--scoreDelOpen −1 -- scoreDelBase −1--scoreInsOpen −1 --scoreInsBase −1 --seedSearchStartLmax 25 -- winAnchorMultimapNmax 100”. Duplicate reads were removed with PICARD (2.9.0) (http://broadinstitute.github.io/picard). Note that duplicate removing is usually not performed for RNA-Seq analysis, but considering that we were looking for extreme gene expression, we wanted to make sure the signals found were not due to duplicates. Fragments mapped to the genes annotated by Ensembl (Version 104) were counted using featureCounts (2.0.3) (48).

### Assignment of extreme gene expression

For a specific gene in a specific sample population, we required that at least one TPM value (CPM value for Drosophila, and mean(CPTT) value for human snRNA-Seq) is larger than 5. If a TPM value is larger than Q3 + 5 × IQR and larger than 5, it is assigned to be an extreme over outlier (OO), and if a TPM value is smaller than Q1 – 5 × IQR, it is assigned to be an extreme under outlier (UO).

For sample populations containing two sexes, such as the sample populations of MUS, SPR, SPI, C57BL/6, mouse pedigree, and human, we additionally performed Fisher’s exact test of independence between outlier assignment and sex for each gene. When the P-Value is smaller than the marginal significance cutoff 0.1, we removed the gene from the extreme outlier gene set, because its expression might just be sex-biased instead of extreme in a few samples. For human sample populations, unlike mouse and fruit fly sample populations which are well controlled during animal experiments in the lab, besides sex, age and death type also introduce significant heterogeneity. Thus, we also performed Fisher’s exact test of independence between outlier assignment and age / death type separately for each gene, and removed the gene from the extreme outlier gene set if any P-Value is smaller than 0.1.

Individual checking of read coverage in annotated non-coding genes showed a number of cases where outlier read mapping covered only a small portion of the genes. While this is in itself an interesting observation, we did not strive to further explore it here. Hence, all the analysis was restricted to protein-coding genes only, which represent roughly 2/3 of the outlier calls among all transcripts.

If a gene and its paralog have high sequence similarity, read alignment might be problematic. Thus, we filtered out genes having at least one paralog with identity larger than 90% calculated by Ensembl Compara (46). This cutoff might be too stringent, i.e., alignment tools, such as STAR, can also perform well on genes with high sequence similarities. But we wanted to make sure the signals found were not due to this problem.

### Subsampling analysis

To evaluate the effects on the cutoff (fold of IQR) and sample size for outlier calling, we used subsampling analysis from the mouse dataset of the 48 DOM individuals. We first excluded one sample from each of the five organs, which contains a very large number of outlier genes and could lead to an uninformative large variance during subsampling. The samples excluded are: H35180 (individual 38) for brain, H35232 (individual 18) for heart, H35342 (individual 35) for kidney, H35302 (individual 31) for liver, and H35425 (individual 30) for mammary gland. For cutoff evaluation, we tested fold of IQR from 3 to 10. For each organ and each fold of IQR, we did 1000 subsampling runs. In each run, we randomly chose 24 samples from the 47 samples in total, and performed the assignment analysis of extreme gene expression as described above. For sample size evaluation, we used 5 as the fold of IQR, and tested the sample size N as 8, 16, 24, 32, and 40. For each organ and each sample size, we also did 1000 subsampling runs. In each run, we randomly chose N samples from the 47 samples in total, and performed the assignment analysis of extreme gene expression as described above.

### Human snRNA-Seq analysis

For the assignment of extreme gene expression in each cell type of the human snRNA-Seq data, we used the mean of the counts per 10,000 reads, mean(CPTT), of all the cells belonging to the cell type in an individual as the expression level of a gene, and then performed the same analysis described above.

For the analysis of the expression variance of OOs, we used the counts per 10,000 reads, CPTT, as the expression level of a gene in a cell. For each of the 307 OOs belonging to the 145 genes, we calculated coefficient of variation (CV) of the gene among the cells of the cell type and the individual which the OO exists. In addition, we also chose a control gene for the OO in the same cell type and the same individual. The rule of the choice is that the mean(CPTT) of the control gene of the individual is within the Q1 to Q3 range across individuals of the cell type, and the mean(CPTT) of the control gene is close to the mean(CPTT) of the OO. Then, we also calculated the CV of the control gene in the same way.

### Human protein data analysis

Blood samples from the NHS and NHSII were analyzed for Prolactin using microparticle enzyme immunoassays in 11 batches using the ARCHITECT chemiluminescence immunoassay system (Abbott Diagnostics), and an additional three batches were assayed using the IMx System (Abbott Laboratory) (26). GH was assayed by ELISA using reagents from Diagnostic Systems Laboratory (27).

## Declarations

### Ethics and permissions

All mice were kept according to FELASA (Federation of European Laboratory Animal Science Association) guidelines, with the permit from the Veterinäramt Kreis Plön: PLÖ-004697. The Government of Schleswig-Holstein provided permission to sacrifice animals under permit number V312-72241.123-34.

The study protocol for the human hormone data analysis was approved by the institutional review boards of the Brigham and Women’s Hospital, Harvard T.H. Chan School of Public Health, and those of participating registries as required.

### Availability of data

The ENA BioProject accession numbers for the sequencing data reported in this study are: PRJEB36991 and PRJEB50011 for the wild mouse populations, PRJEB75700 for the C57BL/6 inbred mouse strain, and PRJEB75139 for the wild mouse families. The supplementary data tables are accessible at Edmond (repository of the MPG) through the private link for reviewers: https://edmond.mpg.de/privateurl.xhtml?token=f3398691-6c5e-4e57-8249-ae5f9deb285e

### Code availability

The code for our outlier detection is at “https://github.com/cxiepku/outlier”.

## Funding

The study was funded by institutional resources of the MPG. We would like to thank the participants and staff of the NHS and NHSII for their valuable contributions as well as the following state cancer registries for their help: AL, AZ, AR, CA, CO, CT, DE, FL, GA, ID, IL, IN, IA, KY, LA, ME, MD, MA, MI, NE, NH, NJ, NY, NC, ND, OH, OK, OR, PA, RI, SC, TN, TX, VA, WA, WY. We thank the Channing Division of Network Medicine, Department of Medicine, Brigham and Women’s Hospital as home of the Nurses’ Health Studies. The authors assume full responsibility for analyses and interpretation of these data. The content is solely the responsibility of the authors and does not necessarily represent the official views of the National Institutes of Health. We would like to acknowledge the following grants: UM1 CA186107, P01 CA87969 (NHS) and U01 CA176726 (NHSII), R01 CA67262 (NHSII) from the National Cancer Institute of the National Institutes of Health, National Natural Science Foundation of China (grant number: 32370665), Guangdong Basic and Applied Basic Research Foundation (grant number: 2024A1515030117).

### Author contributions

C. X. and D. T. have devised the study, C.X. and S.K. have generated transcriptome data, C.X., W.Z. and D.T. have analyzed the transcriptome data, C. H. and S. T. have contributed the human hormone gene analysis, C. X. and D.T. have written the manuscript, with input from all co-authors.

## Acknowledgements

We thank Luisa Pallares for providing *D. melanogaster* data and discussion, Christine Pfeifle and Heike Harre for animal care taking and organ preparation, Michaela Schwarz for RNA purification, Derk Wachsmuth and Kristian Ullrich for IT support.

